## Supplementary Material for "Q4ddPCR (May the Fourth Be Precise): A Flexible, 4-Target Assay for High-Resolution HIV Reservoir Profiling"

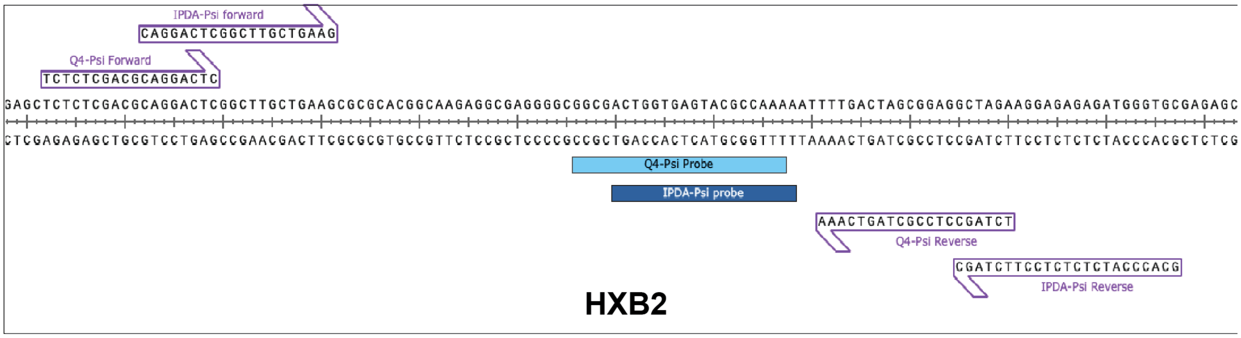

**Supplementary Figure 1 | *Ψ*-binding site of Q4ddPCR**

Primer and probe sequences for the Q4- and IPDA-based *Ψ*-targets mapped to the HXB2 reference genome.

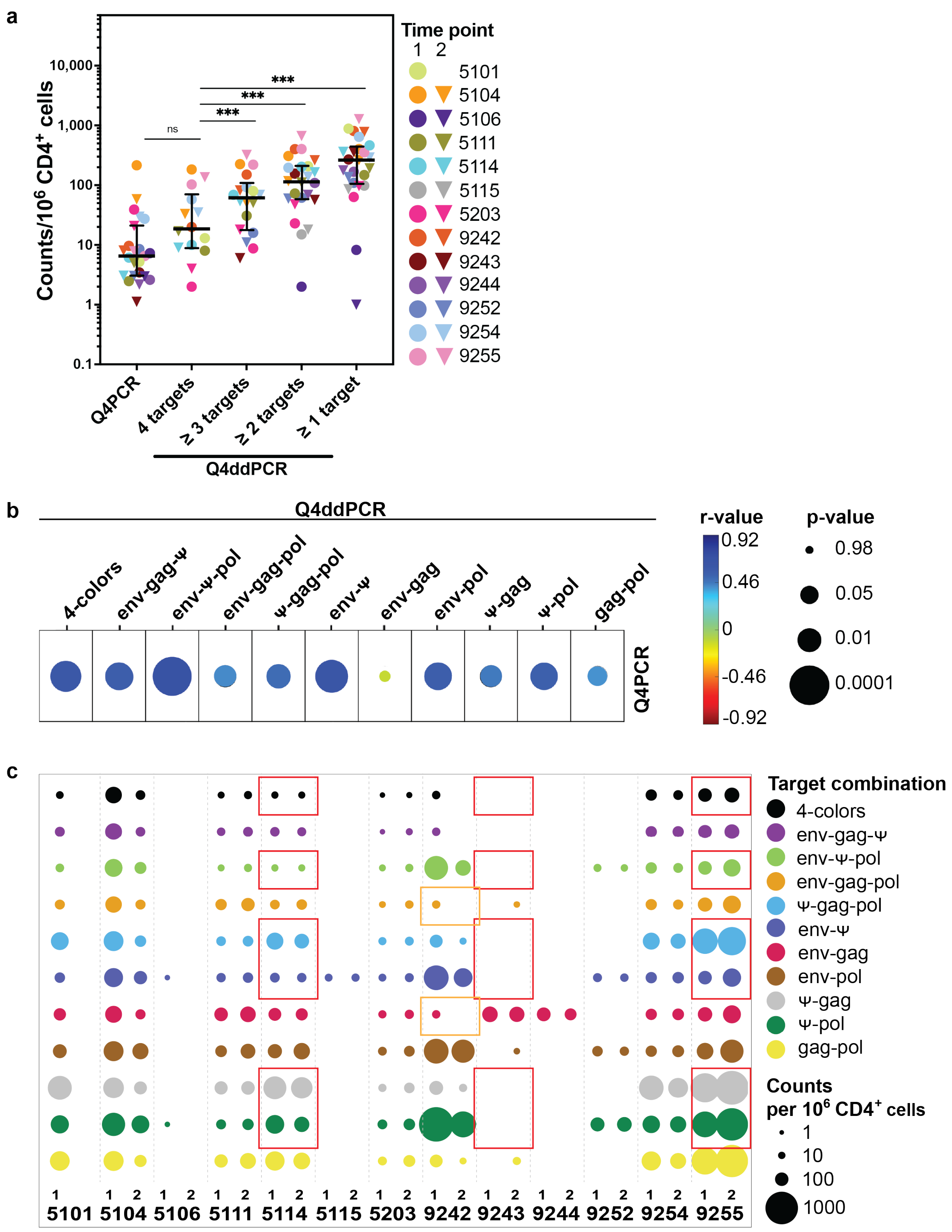
a
Counts/10^6 CD4+ cells
0.1
1
10
100
1,000
10,000
ns
***
***
***
Q4PCR
4 targets
≥ 3 targets
≥ 2 targets
≥ 1 target
Q4ddPCR
Time point
1 2
5101
5104
5106
5111
5114
5115
5203
9242
9243
9244
9252
9254
9255
b
Q4ddPCR
4-colors
env-gag-Ψ
env-Ψ-pol
env-gag-pol
Ψ-gag-pol
env-Ψ
env-gag
env-pol
Ψ-gag
Ψ-pol
gag-pol
Q4PCR
r-value
0.92
0.46
0
-0.46
-0.92
p-value
0.98
0.05
0.01
0.0001
c
Target combination
4-colors
env-gag-Ψ
env-Ψ-pol
env-gag-pol
Ψ-gag-pol
env-Ψ
env-gag
env-pol
Ψ-gag
Ψ-pol
gag-pol
Counts
per 10^6 CD4+ cells
1
10
100
1000
5101 5104 5106 5111 5114 5115 5203 9242 9243 9244 9252 9254 9255

**Supplementary Figure 2 | Validation of Q4ddPCR on longitudinal samples from 13 people with HIV (PWH).**

Q4ddPCR was applied to longitudinal samples from 13 PWH previously characterized using Q4PCR. Q4PCR combines a 4-target qPCR with near full-length genome sequencing and shares the *env, gag* and *pol*-primer/probe sequences with IPDA-based Q4ddPCR and all primer/probe sequences with Q4-based Q4ddPCR. All samples positive for ≥2 targets by Q4PCR had previously been sequenced, resulting in 3,650 proviral sequences for these 13 PWH.

**a** Proviral counts per $10^6$ $CD4^+$ T cells measured by Q4ddPCR across different readout combinations, compared to intact proviral counts from Q4PCR. Cumulative counts from Q4-based Q4ddPCR are grouped by the number of detected targets (1-4). Each symbol represents an individual sample; circles indicate the first time point, triangles the second. Samples from the same participant are color-matched. Medians and interquartile ranges are shown. Statistical comparisons used the two-sided Wilcoxon signed-rank test with post hoc correction for multiple comparisons. ns, not significant; *$p < 0.05$; **$p < 0.01$; ***$p < 0.001$; ****$p < 0.0001$.

**b** Spearman correlation between IPDA-based Q4ddPCR readouts and reservoir size as previously measured by Q4PCR. Correlations are shown for different Q4ddPCR target combinations (columns) with intact proviruses per $10^6$ $CD4^+$ T cells measured by Q4PCR. Spearman's *r*-values are color-coded (blue to red); circle size indicates *p*-value.

**c** Frequency of proviruses positive for various 2-, 3-, or 4-target combinations per $10^6$ $CD4^+$ T cells across two time points in 13 PWH. Each circle represents one readout; size indicates abundance of each target combination; color denotes the specific combination. Participant IDs are shown along the x-axis; samples from two time points are labeled (1, 2), except for participant 5101 (single time point available). Red squares highlight samples with *Ψ* detection failure in one Q4ddPCR variant that was rescued by the alternate version (Supplementary Fig. 2). Orange squares highlight sequence-confirmed intact, but *env*-*Ψ*-negative proviruses from participant 9242 that declined over time. IPDA-based Q4ddPCR results are shown.

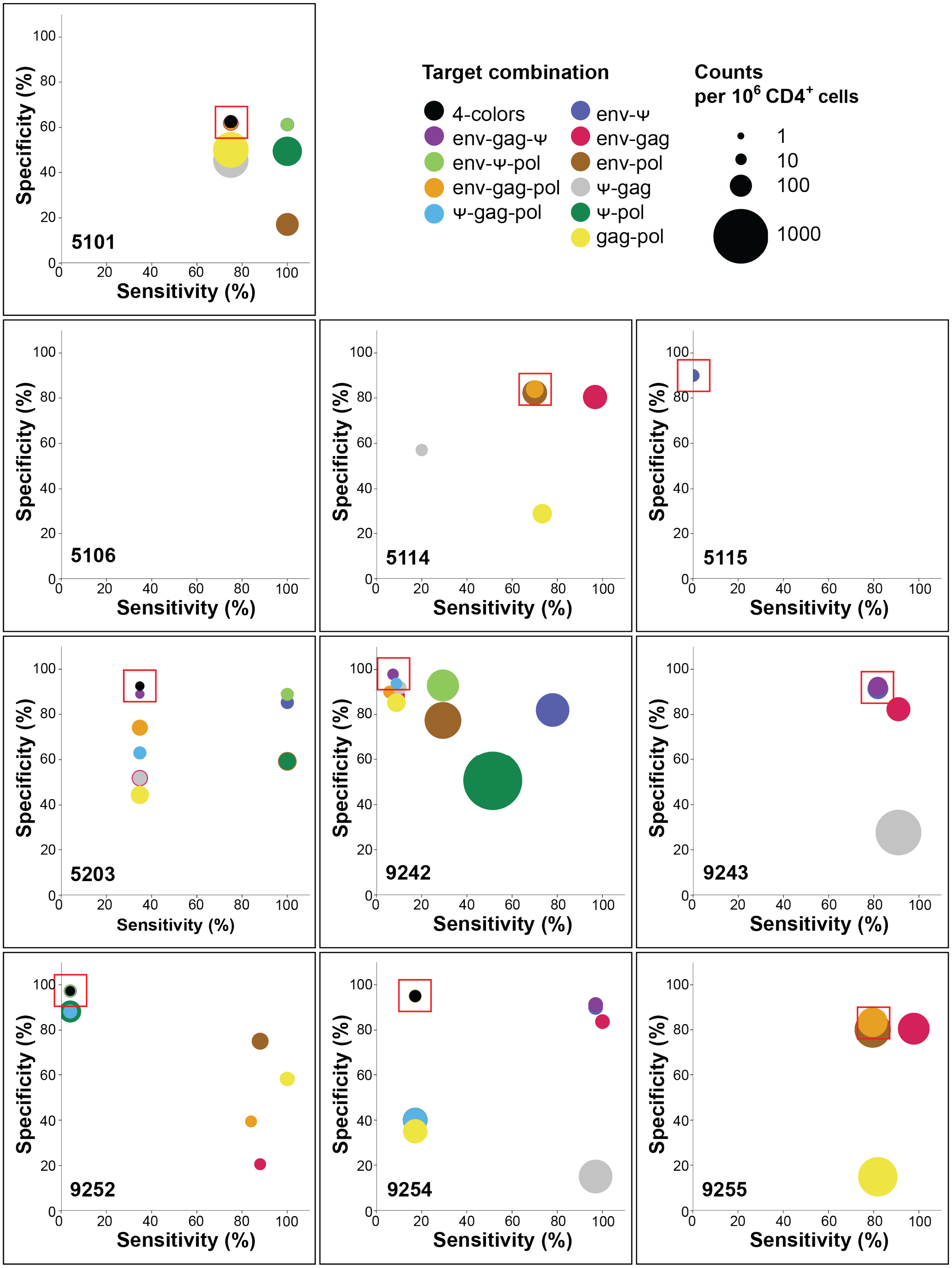

Target combination
4-colors
env-gag-Ψ
env-Ψ-pol
env-gag-pol
Ψ-gag-pol
env-Ψ
env-gag
env-pol
Ψ-gag
Ψ-pol
gag-pol
Counts per 10⁶ CD4⁺ cells
1
10
100
1000
5101
5106
5114
5115
5203
9242
9243
9252
9254
9255
Specificity (%)
Sensitivity (%)

**Supplementary Figure 3 | Sensitivity and specificity of Q4ddPCR readouts for detecting intact HIV proviruses.**

Sensitivity and specificity for intact provirus detection were evaluated across distinct Q4ddPCR target combinations using participant-matched Q4PCR-derived near full-length proviral sequences as a reference across 13 people with HIV. Sensitivity was calculated as the number of sequence-confirmed intact proviruses detected by a given target combination divided by the total number of intact sequences. Specificity was defined as the fraction of defective sequences not detected by the same target combination, relative to all defective sequences. Primer and probe sequences of Q4-based Q4ddPCR match those from Q4PCR. Panels show data from Q4-based Q4ddPCR results of individual participants. Circles denote specific target combinations; size corresponds to the number of proviruses detected per $10^6$ $CD4^+$ T cells, and color encodes specific combination of amplified targets. Red squares mark the target combination selected by the decision tree.

| **Supplementary Table 1 \| Cohort Characteristics** | | | | |
|---|---|---|---|---|
| **Participant ID** | 5101 | 5104 | 5106 | 5111 |
| **Sex** | male | male | male | male |
| **Age at baseline (years)** | 52 | 35 | 31 | 55 |
| **Time on ART at baseline (years)** | 12 | 7 | 6 | 16 |

**Supplementary Table 1| Cohort Characteristics**

All participants were enrolled in studies on broadly neutralizing antibodies and had undetectable viral load at sampling. While all were on suppressive ART at the first sampling time point, treatment was interrupted at the second time point as part of an analytical treatment interruption.

| Supplementary Table 2 \| Sensitivity and Specificity of Specific Target Combinations | | | | |
|---|---|---|---|---|
| Q4PCR Target Combination | Sensitivty (%) | Specificity (%) | Positive Predictive Value (%) | Negative Predictive Value (%) |
| ≥ 1 target | 82.3 | 41.8 | 20.3 | 96.4 |
| ≥ 2 targets | 67.3 | 69.7 | 28.6 | 93.7 |
| ≥ 3 targets | 55 | 86.6 | 42.5 | 91.7 |
| 4-targets | 45.5 | 95.3 | 63.5 | 90.3 |
| env | 93.2 | 45.3 | 23.5 | 98.5 |
| Ψ | 79.2 | 63.9 | 28.4 | 95.8 |
| gag | 87.3 | 26.7 | 17.7 | 97.3 |
| pol | 69.4 | 31.3 | 15.4 | 94.1 |
| env-Ψ | 73.3 | 91.1 | 59.7 | 94.7 |
| env-gag | 81.7 | 63.7 | 28.9 | 96.3 |
| env-pol | 64.9 | 65.5 | 25.3 | 93.3 |
| Ψ-gag | 64.3 | 75 | 31.7 | 93.2 |
| Ψ-pol | 57 | 77.6 | 31.4 | 92 |
| gag-pol | 62.4 | 45.4 | 17.1 | 92.9 |
| env-gag-Ψ | 61.8 | 94.5 | 66.9 | 92.8 |
| env-Ψ-pol | 51.3 | 94.1 | 61.1 | 91.2 |
| env-gag-pol | 59.1 | 72 | 27.6 | 92.4 |
| Ψ-gag-pol | 47.8 | 85.6 | 37.5 | 90.7 |
| 4-color | 45.5 | 95.3 | 63.5 | 90.3 |

**Supplementary Table 2**
